# SNP Selection and Concordance in Consumer Genetics Testing

**DOI:** 10.1101/352732

**Authors:** John F. Thompson

## Abstract

The use of Direct To Consumer (DTC) genetic testing for predicting health risks and a variety of other phenotypes has been extensively discussed. Additionally, there have been wide ranging discourses on privacy and ethical concerns. Much less attention has been paid to what most people actually use DTC testing for: ancestry determination. Furthermore, comparison of the platforms used by different companies and how they have chosen SNPs to address the questions of health and ancestry have not been broadly reported. When SNPs across three genotyping platforms are compared, only 16-18% of SNPs with reported genotypes are shared across all platforms. Only 110,051 of the more than 600,000 SNPs are called on all three panels examined (Ancestry, 23andMe and MyHeritage). SNPs genotyped on all platforms are highly concordant with only two SNPs having discordant calls. When the SNPs unique to a single panel are examined, it is apparent that each company has its own strategy for choosing SNPs. When each platform is examined, the unique SNPs have different frequencies, ethnic selectivities, and chromosomal locations. Because each company separates the world into different, overlapping geographical regions, it is impossible to do an exact comparison of ancestry results. Factoring in the ways the regions overlap, congruent results are generated for the major contributors to ancestry.

## Introduction

The recent approval of 23andMe′s test for three BRCA1/BRCA2 mutations (www.fda.gov/NewsEvents/Newsroom/PressAnnouncements/ucm599560.htm) has sparked renewed discussion of the impact of such tests on individual health. The benefits of knowing whether one has a pathogenic variant that predisposes to disease is countered by the disadvantages of having a potentially inaccurate test and/or a lack of understanding of the test’s meaning by the individual ordering it. These issues have been extensively discussed elsewhere (1–8) and will not be addressed here in detail. While many people are interested in the health information that can be potentially gleaned from the genome-wide SNP data, most actually purchase the test for its use in helping to understand family history. Even the names of many DTC companies (Ancestry, MyHeritage, FamilyTreeDNA) reinforce the idea that the primary use for the data is family history. However, the issues raised by the health tests have diminished the attention paid to how the family history data is actually generated and the potential differences among providers.

There are three required steps for determining ancestral origins via DNA testing. The first component, algorithms for determining ancestry, has been described at a high level but the methods are mainly proprietary and have sparked patent battles (www.wired.com/story/23andme-sues-ancestry/). As a result, the methods cannot be readily compared. The second component, the reference populations, are similarly proprietary. Each company has its own set of samples representing populations around the world. The ability of these samples to accurately reflect a particular area cannot be determined without access to them and how they were selected. The third component, the SNPs used for the analysis, however, can be compared across platforms. The SNPs chosen for the three panels are quite dissimilar so can be contrasted to determine differences in strategy based on publically available information.

## Methods

Ancestry, 23andMe, and MyHeritage all offer similar DTC tests that employ the Illumina Infinium microarray (https://www.illumina.com/products/by-type/microarray-kits/infinium-iselect-custom-genotyping.html) that provides genotyping data for a custom list of around 700,000 SNPs. Each company chooses its own set of SNPs for genotyping. When data is provided back to the consumer, it includes both chromosomal locations and dbSNP numbers when available, allowing easy comparison of SNPs and genotype calls. In instances where there are no dbSNP identifiers, chromosomal locations were provided. In addition to comparing genotype files, databases used to obtain information were dbSNP (www.ncbi.nlm.nih.gov/projects/SNP/). the hg19 human reference genome (www.ncbi.nlm.nih.gov/assembly/GCF_000001405.13/?report=full), the gnomAD sequence database (gnomad.broadinstitute.org/) and the BLAT tool for comparing short DNA sequences (http://genome.ucsc.edu/cgi-bin/hgBlat?command=start).

## Results

Three vendors with similar products, Ancestry, 23andMe, and MyHeritage were chosen to perform genotyping and interpretation. The MyHeritage dataset was also sent to two other vendors, FamilyTreeDNA and Gencove, for them to interpret independently. Each platform reports over 600,000 SNP genotypes so SNPs are located about every 5000 bp. Because accurate SNP genotyping requires well-behaved DNA, an even distribution of SNPs across the whole genome is not possible. If DNA is too extreme in GC-content, either high or low, discrimination of SNPs is not possible. Also, if a SNP is embedded in a region that is too similar to other genomic regions, the signal from the different regions will interfere with each other, leading to potentially erroneous calls. Despite sequence limitations, there are no gaps between SNPs >50 Mb with any vendor. 23andMe has the lowest number of gaps >1 Mb (19) while MyHeritage has the highest (27).

As shown in Table 1, the chromosomal SNP distributions are very similar for chr1-22. However, SNP densities on the mitochondrial and sex chromosomes are quite different between vendors. MyHeritage and 23andMe have similar numbers of SNPs on chrX (2.5 and 2.6%) while Ancestry has 4.2% on the non-recombining regions of chrX (which they refer to as chr23) and another 0.3% on the pseudo-autosomal regions of chrX (referred to as chr24). These regions of chrX have high homology to chrY. MyHeritage and Ancestry have similar numbers of SNPs on chrY (Ancestry′s chr25) while 23andMe has more than 7 times as many SNPs. MyHeritage does not report any mitochondrial SNPs while Ancestry reports 164 (chr26) and 23andMe reports 4301. Each of these chromosomes (X, Y, mito) has unique attributes with respect to determining ancestry along maternal and paternal lines.

**Table 1.**
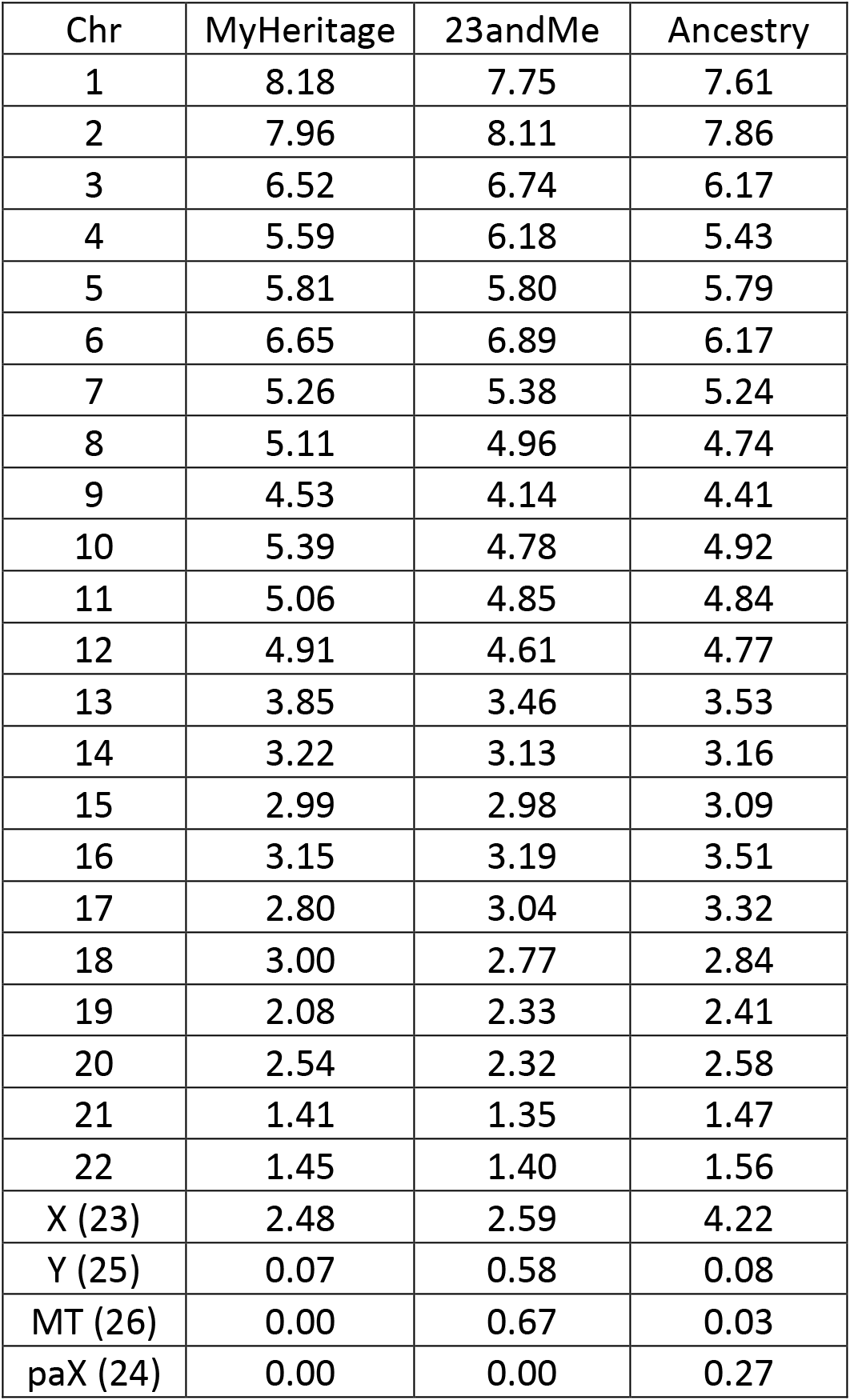
Per cent of total SNPs on each chromosome

| Chr | MyHeritage | 23andMe | Ancestry |
| --- | --- | --- | --- |
| 1 | 8.18 | 7.75 | 7.61 |
| 2 | 7.96 | 8.11 | 7.86 |
| 3 | 6.52 | 6.74 | 6.17 |
| 4 | 5.59 | 6.18 | 5.43 |
| 5 | 5.81 | 5.80 | 5.79 |
| 6 | 6.65 | 6.89 | 6.17 |
| 7 | 5.26 | 5.38 | 5.24 |
| 8 | 5.11 | 4.96 | 4.74 |
| 9 | 4.53 | 4.14 | 4.41 |
| 10 | 5.39 | 4.78 | 4.92 |
| 11 | 5.06 | 4.85 | 4.84 |
| 12 | 4.91 | 4.61 | 4.77 |
| 13 | 3.85 | 3.46 | 3.53 |
| 14 | 3.22 | 3.13 | 3.16 |
| 15 | 2.99 | 2.98 | 3.09 |
| 16 | 3.15 | 3.19 | 3.51 |
| 17 | 2.80 | 3.04 | 3.32 |
| 18 | 3.00 | 2.77 | 2.84 |
| 19 | 2.08 | 2.33 | 2.41 |
| 20 | 2.54 | 2.32 | 2.58 |
| 21 | 1.41 | 1.35 | 1.47 |
| 22 | 1.45 | 1.40 | 1.56 |
| X (23) | 2.48 | 2.59 | 4.22 |
| Y (25) | 0.07 | 0.58 | 0.08 |
| MT (26) | 0.00 | 0.67 | 0.03 |
| paX (24) | 0.00 | 0.00 | 0.27 |

Each of the vendors deals with reporting ChrX genotypes in a different way. All MyHeritage genotypes are presented as diploid and nearly all those calls are homozygous in this male sample (30 calls across the entire chromosome are reported as heterozygous). 23andMe makes diploid calls in the pseudo-autosomal regions with their boundaries of ChrX:153,977-2,697,868, and ChrX:154,939,018- 155,234,707. Haploid calls are made in the vast majority of the chromosome (2,700,163-154,844,425). Ancestry splits ChrX into two separate chromosomes, the bulk of chrX (called Chr23) and the pseudo-autosomal regions referred to as Chr24. The boundaries of Chr23 and Chr24 overlap significantly with Chr23 spanning 2,700,157-154,929,412 and Chr24 spanning 2,655,180-28,817,458. No SNPs in the other region generally assigned as pseudo-autosomal are listed (9). In theory, there should be no heterozygous SNPs in the non-recombining part of chrX in a male. However, Ancestry reports 57 heterozygous SNPs on chr23. Of these 57 SNPs, 15 are also reported by MyHeritage and 8 by 23andMe. All of the 23andMe SNPs were reported as ″no call″ while 11/15 MyHeritage SNPs also were reported as heterozygous with 4 ″no calls″. If the sequence surrounding these SNPs is examined and compared to the rest of the genome using BLAT, 53/57 have sequences on other chromosomes that are sufficiently similar to the region around the chrX SNP that it could interfere with the call. 29 of these similar sequences are only on chrY while 24 have similarities on chrY as well as other chromosomes including some with potentially dozens of interfering sequences (rs2574221, rs5983978, rs5984925). Because genotyping technology is tuned to use allele ratios for determining calls and is less sensitive to absolute signal, the presence of additional homologous sequences can lead to erroneous results because the chrX signal cannot be distinguished from chrY and other sequences.

It has been reported that 40% of DTC SNP results are incorrect when replication is attempted via sequencing in a clinical laboratory (10). Such a high error rate would obviously lead to ancestry errors as well. To determine the concordance of SNPs within this sample set, all 110,051 SNPs on autosomal chromosomes (1-22) with genotypes reported on all three platforms were compared. Only four genotypes were discordant. However, inspection of the genotypes revealed that two apparent SNP discordances (rs12721636, rs28365067) were the result of faulty nomenclature where the wrong strand was reported in the genotype so the calls were, in actuality, concordant. The two other discordances, rs9639507 and rs34923550, had different calls (heterozygous vs. homozygous) so were, in fact, technical errors. There were no obvious sequence features in these SNP regions that contributed to these discordances. The two discordances among the 110,051 SNPs leads to concordance of >99.999%. Typically, such arrays are found to be ~99.8% accurate (11) but this set of 110,051 is likely heavily biased towards well-behaved SNPs by selection and retention by three separate vendors.

While the 110,051 SNPs common to all platforms provide uniform information, there are another ~500,000 SNPs on each platform that are not shared by one or both other platforms. To better understand, how each company chooses SNPs, autosomal SNPs unique to a single platform were examined and compared to each other and the universally shared SNPs. As shown in Table 2, shared SNPs have a much higher rate of heterozygosity, indicating a much higher average minor allele frequency. Unique SNPs on Ancestry and MyHeritage have similar heterozygosity rates compared to each other while 23andMe is significantly lower, with a heterozygosity rate less than 1/3 of the shared SNPs, indicating that most unique SNPs have a much lower variant frequency.

**Table 2.** Autosomal SNP calls

| Vendor | Homozygous Calls | Heterozygous calls | Heterozygous % |
| --- | --- | --- | --- |
| Ancestry unique | 150,820 | 38,953 | 20.5 |
| 23andMe unique | 350,458 | 42,654 | 10.9 |
| MyHeritage unique | 181,508 | 50,549 | 21.8 |
| Shared | 68,221 | 37,182 | 35.3 |

To look at representative SNPs in more detail, all unique SNPs were sorted by chromosomal position and 100 SNP trios were randomly picked that included one SNP from each vendor in consecutive order on the genome. Each trio was further filtered such that the three SNPs were within 5000 bp of each other, all had genotype calls, and all were found in the gnomAD sequence variant database (gnomad.broadinstitute.org/). Each set of three SNPs was selected to be separated by >10Mb from all other trios examined in order to ensure independence of the trios. While each vendor′s reference populations undoubtedly had a major impact on their SNP choices, these databases are not publicly available for evaluation. Instead, it is necessary to use public databases as surrogates to evaluate relevant SNP properties. The gnomAD database provides good data for variant frequency and ethnic specificity (12). Each SNP has been ″genotyped″ (actually sequenced) in about 30,000 individuals if the SNP is in non-exonic regions and in about 270,000 individuals if the SNP occurs in an exonic region. These individuals are broken down into at least 6 different ancestral groups (African, Ashkenazi Jewish, East Asian, European except Finland, Finnish, Latino, Other, and, with exonic SNPs, South Asian). These data provide an accurate overview of SNP frequencies over continental regions though the database may miss high frequency SNPs within more localized geographical regions that may be important for the vendors.

To compare SNP choices, SNPs were categorized in two ways, by overall frequency (less than 1%, 1-5%, 5-20% and greater than 20%) for the minor allele (which was sometimes the reference allele) and also by the difference in frequency between the highest frequency population and the lowest frequency population (less than 2-fold, 2-5 fold and greater than 5-fold). The SNPs for each vendor differed in how these categories were populated. The most common category for Ancestry was greater than 20% minor allele frequency and 2-5-fold difference between populations. The most common category for MyHeritage was 5-20% minor allele frequency and >5-fold discrimination. The most common category for 23andMe was the high population difference category (>5-fold) but lower overall frequency (1-5%). All vendors chose at least 50% SNPs with high population differences but this choice was most extreme with 23andMe (79%). This was at the expense of picking very low frequency SNPs (32% less than 1% frequency) while Ancestry and MyHeritage chose far fewer SNPs in that frequency range (8% and 3%, respectively). This fits well with the heterozygosity rates observed in the DNA analyzed here because the less common SNPs will frequently be homozygous reference.

**Table 3.** Unique SNP characteristics

| Ancestry | Ethnicity Ratios |  |  | 23andMe | Ethnicity Ratios |  |  | MyHeritage | Ethnicity Ratios |  |  |
| --- | --- | --- | --- | --- | --- | --- | --- | --- | --- | --- | --- |
| MAF | <2 | 2-5 | >5 | MAF | <2 | 2-5 | >5 | MAF | <2 | 2-5 | >5 |
| <1% |  |  | 8 | <1% | 1 |  | 31 | <1% |  |  | 3 |
| 1-5% |  |  | 13 | 1-5% |  |  | 40 | 1-5% |  |  | 23 |
| 5-20% |  | 8 | 22 | 5-20% | 1 | 11 | 7 | 5-20% | 2 | 13 | 25 |
| >20% | 8 | 34 | 7 | >20% | 4 | 4 | 1 | >20% | 8 | 20 | 6 |
| Total | 8 | 42 | 50 | Total | 6 | 15 | 79 | Total | 10 | 33 | 57 |

Comparison of SNP properties is relatively straightforward because of the common nomenclature and public data. Comparison of ancestry determination is more problematic because each vendor has separated the world into different geographical sectors. Thus, it is generally impossible to directly compare the detailed results at a country-level resolution. An attempt has been made to make the regional definitions equivalent in Table 4 but some quantitative variation is undoubtedly due to different borders for the locations of ancestral reference populations.

**Table 4.**
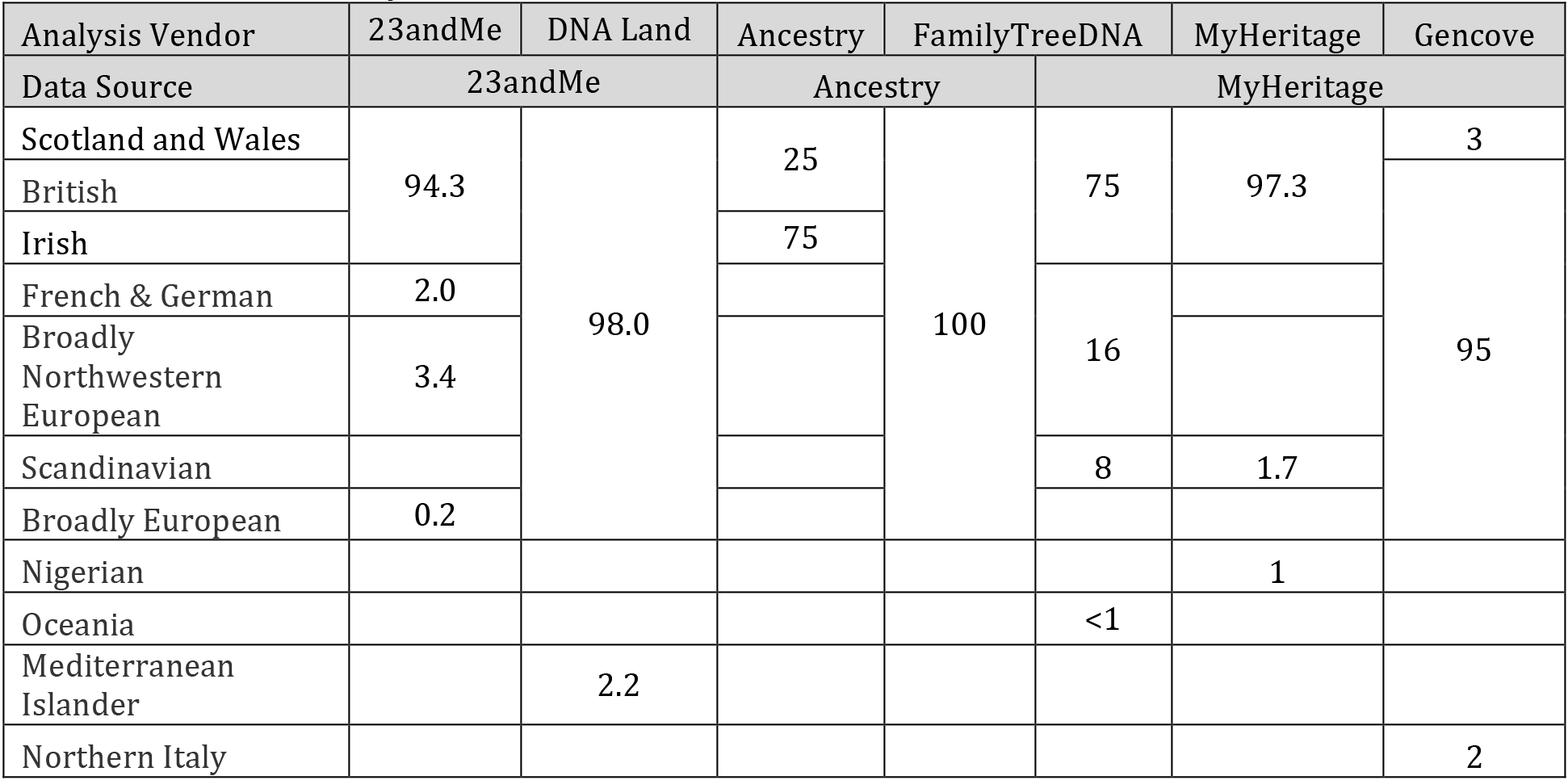
Ancestral analysis

The major European contributions to ancestry match the documentary evidence to the extent it is known (75% Irish, 19% English, 3% Scottish, 3% Swedish). Some of the minor contributions vary across analyses with less than 3% attributed to a variety of sources, Mediterranean Islander, Oceania, Nigeria, and Northern Italy. While any of these are possible, the low levels and inconsistencies across analyses suggest they are more likely noise and not real.

## Discussion

Most discussions of DTC genetic testing have focused on disease diagnosis or ethical/privacy issues. Only rarely has there been a discussion of the data quality and its use for ancestry testing. Previous reports that compared genotyping results examined panels of very different sizes (13, 14) so the comparison was necessarily limited in scope. While they found high concordance, there were too few SNPs compared to draw strong conclusions. One widely reported study indicated that ″40% of variants in a variety of genes reported in DTC raw data were false positives″ (10), suggesting a very different technical quality. The high degree of technical reproducibility we found stands in stark contrast to this result. Only 2 of the 110,051 shared SNPs had a single discordant call among them. This high level of accuracy, greater than 99.999%, is seemingly at odds with the other study.

However, knowing how the technology works explains the apparent discrepancy between the generally very high accuracy demonstrated here and the high rate of errors among disease variant calls. The genotyping technology employed by all vendors works best with common variants where there are individuals with both reference and alternate homozygous alleles as well as individuals with heterozygous alleles. In such a situation, the boundaries between the calls are more distinct and borderline calls can be more easily distinguished and discarded. However, the disease-causing variants sent for clinical confirmation are rare and thus much more likely to be inaccurate than common SNPs. This highlights the difficulties in using DTC SNP genotyping for analyzing rare disease variants and reinforces the need to use proper, clinically-validated tests for disease diagnosis or confirmation of DTC findings if potential issues are identified via DTC testing.

The DNA-based ancestry analyses using six companies to provide results using three sets of data yield similar conclusions. The primary problem in making a quantitative comparison among vendors lies in the fuzziness of the geographical boundaries that each provides. Not surprisingly, the reference populations originate from large areas with borders that have varied over time. Even if the birth location for current DNA donors is known precisely, the birth locations for their ancestors will be known with less certainty. As a result, each company defines the borders for their analysis differently.

With these data, all vendors agree the DNA is of primarily Irish/English/Northwestern European origin. This DNA-derived ancestry is consistent with documentary evidence obtained independently. The major differences among vendors are contributions of less than 3%. This could be simply noise, errors in genotyping/haplotyping, less than perfect reference populations or may actually reflect real contributions from distant, undocumented ancestors. Higher resolution studies and/or better reference populations would be needed to clarify these discrepancies.

Using all three vendors for genotyping and three more for interpretation provided more confidence than simply using a single vendor would. It is easier to see where the uncertainties are with both percentages and geography. It supports documentary evidence of approximately 75% Irish, 19% English, 3% Scottish, and 3% Swedish. There is still some space for uncertainty at the <3% level that will require better tests and reference populations to resolve.

